# Elucidating the molecular landscape of centronuclear myopathies in mouse models: an integrative multi-omics and network analysis

**DOI:** 10.1101/2025.01.03.631202

**Authors:** Alix Simon, Charlotte Gineste, David Reiss, Julie Thompson, Jocelyn Laporte

## Abstract

Centronuclear myopathies (CNM) are rare inherited muscle disorders characterized by muscle atrophy, weakness, and altered muscle fiber structure, primarily due to mutations in genes like MTM1, DNM2, and BIN1. The pathomechanisms implicated in CNM are only partially understood, and no curative therapies are available for patients. This study exploits a unique multi-omics dataset and network-based analyses to elucidate the molecular pathways involved in CNM. First, we performed Weighted Gene Co-expression Network Analysis (WGCNA) to identify gene modules correlated with CNM phenotypes. We find that modules correlated to a positive muscular phenotype were enriched for genes involved in muscle contraction, RNA processes and oxidative phosphorylation, while modules associated with impaired muscle structure and function were enriched in immune response, innervation, vascularization processes, and fatty acid oxidation. Next, we integrated transcriptomic, proteomic, and metabolomic data from the *Mtm1*^-/y^ mouse model with public knowledge bases using a multilayer network approach and explored the network using a random walk with restart approach. This study revealed novel metabolites that could be targeted through dietary supplementation as potential therapeutic strategies. Our findings demonstrate how multi-omics network analyses can reveal new aspects of CNM pathomechanisms and identify avenues for intervention.

## Introduction

Centronuclear myopathies (CNM) are a group of rare inherited muscular disorders characterized by muscle weakness, hypotonia, and abnormal positioning of nuclei within muscle fibers^1^. These diseases are primarily caused by mutations in genes such as *MTM1*, *DNM2*, and *BIN1*. Despite progress in identifying causative mutations, the intricate molecular pathways leading from these genetic disruptions to the clinical manifestation of CNM are still not fully delineated, limiting the development of effective therapies.

Omics approaches, including transcriptomics, proteomics, and metabolomics, offer powerful tools for uncovering the broader molecular landscape involved in CNM. These approaches represent comprehensive and unbiased techniques to characterize complex biological systems, enabling the identification of key molecular mechanisms underlying various processes^2^. Single-omic analysis often involves differential expression analysis between two conditions to identify dysregulated entities. Such analyses can yield valuable insights, but they cannot take into account the intricate regulatory and interaction processes that connect different layers of omics data, and they are limited to pairwise comparisons. Therefore, integration of multi-omics data together with clinical information has shown great promise in achieving a holistic understanding of complex biological systems^3^.

Integration approaches based on biological networks have gained traction in recent years. Weighted Gene Co-expression Network Analysis (WGCNA) is one such approach that can be used to construct networks based on the correlation patterns between genes across multiple samples and to identify gene modules that correlate with clinical traits^4^. Multilayer network approaches, which integrate diverse biological data types (e.g., transcriptomic, proteomic, and metabolomic data), data sources (e.g., protein-protein interactions, pathway co-membership, co-expression), and/or experimental conditions (e.g., transcriptomic data obtained from several models) offer an even more comprehensive framework^5^. By leveraging high-throughput data from multiple layers, these methods allow modeling of complex, multi-dimensional relationships that single-layer network approaches cannot capture.

Here, we used network integration techniques to uncover novel molecular interactions and pathways that contribute to disease pathomechanisms in CNM. First, we performed WGCNA on several cohorts of CNM murine models treated with various therapeutic approaches. Then, we performed a case study focusing on the investigation of the multi-modal pathomechanisms implicated in XLCNM (X-linked CNM, caused by mutations in *MTM1*), for which comprehensive experimental transcriptomics, proteomics, and metabolomics data are available. Thus, we built and explored a heterogeneous multilayer network integrating the multi-omic data generated for the *Mtm1*^-/y^ mouse model with biological knowledge extracted from publicly available databases.

## Results

### Weighted gene correlation network analysis of several models of CNM

#### Transcriptomic and phenotypic data collection

To identify genes implicated in skeletal muscle structure and function and in the pathogenesis of CNM, we performed WGCNA on several murine models of myopathies treated with various therapeutic approaches (Table 1). In particular, three models of CNM (*Mtm1*^-/y^, *Bin1*^mck-/-^, *Dnm2*^S619L/+^) were considered. Therapeutic approaches included gene expression modulation by genetic crossing or antisense oligonucleotide (ASO) and tamoxifen repurposing. For each cohort, bulk RNA-sequencing data was retrieved from previous publications^6,7^ or generated for this study (cohort F). For all cohorts except cohort F, 4 groups were studied: WT, disease model, WT treated, disease model treated. In cohort F, 3 groups were considered: WT, disease, and disease treated. In total, bulk RNA-seq data of 106 mice were used. Overall, Principal Component Analysis (PCA) of normalized and variance stabilized counts revealed a separation of samples according to their status for WT, WT treated, and disease groups along PC1, which explained 22% of the variance (Fig. 1). Notably, tamoxifen-treated WT samples were an exception to this pattern, clustering on the right side of PC1 alongside disease samples. For the disease groups, we observed that *Mtm1*^-/y^ samples (cohorts A-C) clustered together on the top of PC2. Disease treated samples exhibited a variable distribution on the PCA plot, with some clustering alongside WT samples and others aligning with disease samples in a cohort-specific manner. In particular, disease models treated with tamoxifen diet supplementation (cohorts C, F, and G) tended to cluster with disease samples. In contrast, disease-treated samples in other cohorts were positioned closer to WT samples, especially in cohorts B, D, and E.

**Fig. 1.**
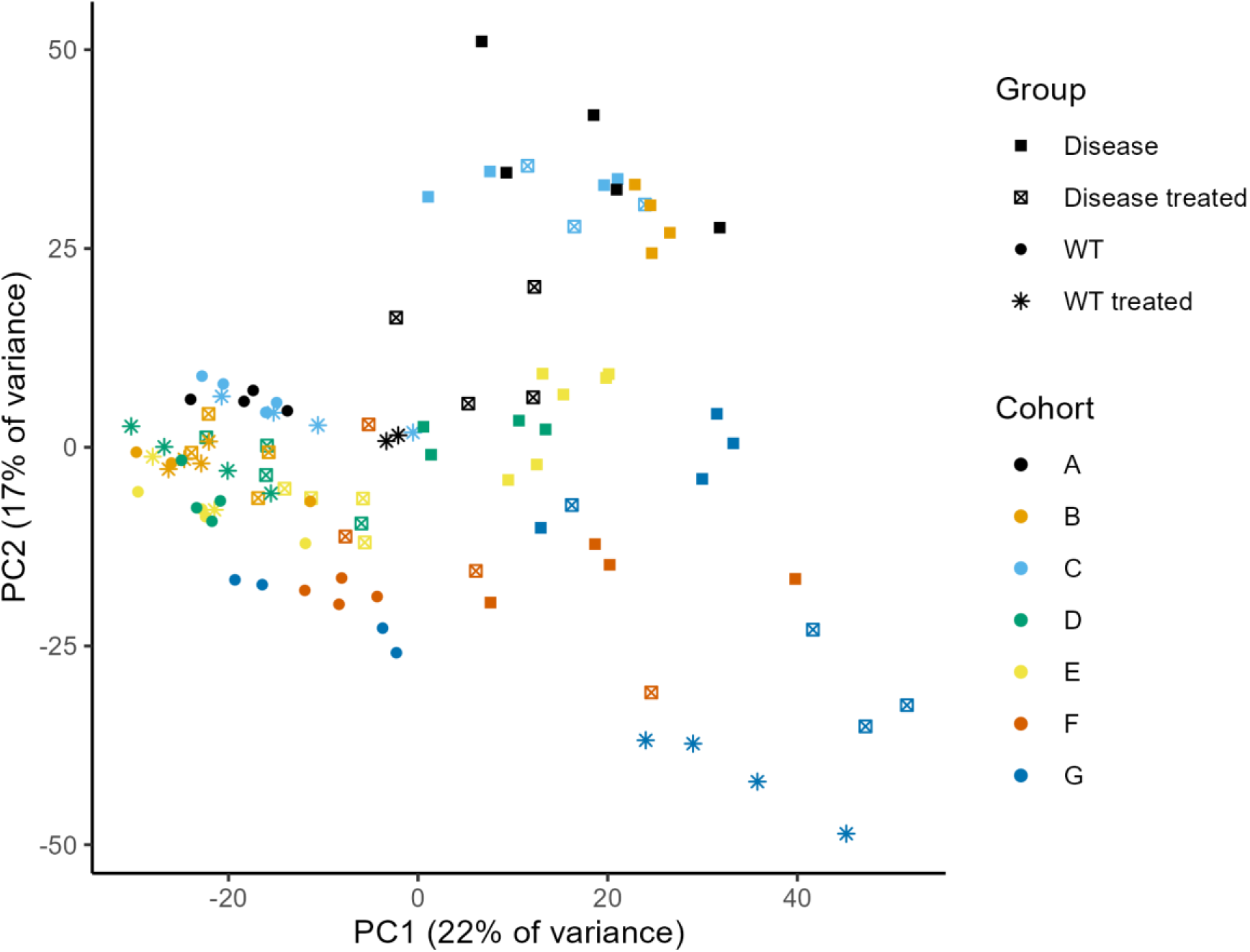
Principal Component Analysis of gene expression in the samples used for WGCNA. Shapes represent the experimental groups, while colors represent cohorts.

**Table 1.**
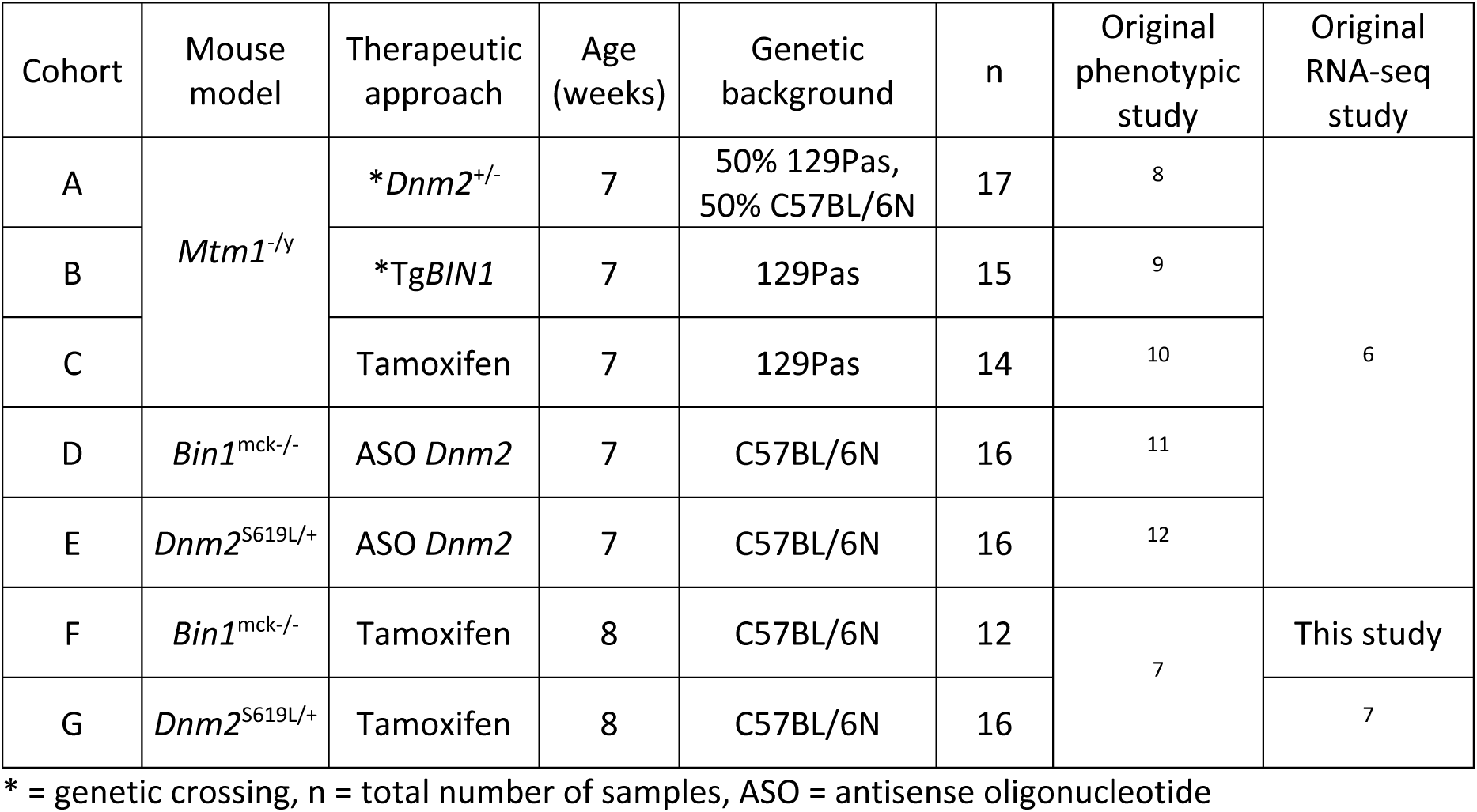
Overview of the transcriptomic cohorts used for WGCNA.

Phenotypic characterizations of the cohorts were retrieved from their respective publications^7–12^. In particular, we retrieved the average value of the following metrics for each group when they were available: body mass (g), tibialis anterior (TA) mass normalized by body mass (mg/g), maximal specific force of the TA relative to WT, hanging test (time to fall off an upside-down wire cage lid) (s), percentage of muscle fibers with internalized or central nuclei, percentage of muscle fibers with abnormal mitochondria localization, percentage of small muscle fibers (fiber diameter < 40 μm). The full phenotypic data is available in Supplementary Information S2. In the absence of mouse identifiers connecting the phenotypic and transcriptomic studies, mice were assigned their average group value for each phenotypic trait.

#### Identification of gene co-expression modules correlated to phenotypic traits

WGCNA was performed to build a gene expression correlation network and identify modules of highly correlated genes. Then, the correlation between module expression and phenotypic traits was computed (Fig. 2). In total, 24 modules were identified, with size ranging from 47 to 1848 genes. Moderate to high correlation coefficients (absolute values ranging from 0.3 to 0.7) were observed for some modules and traits, with the strongest positive and negative correlations observed for normalized TA mass. Module-trait correlation showed low specificity, as most modules were significantly correlated with multiple phenotypic traits, suggesting potential redundancy among these traits. We defined beneficial modules as modules that were positively correlated to normalized TA mass, relative maximal force, and/or hanging time, and negatively correlated to the percentage of fibers exhibiting central nuclei, abnormal SDH staining and/or a small diameter. Modules that showed an opposite correlation pattern compared to beneficial modules were defined as pathogenic.

**Fig. 2.**
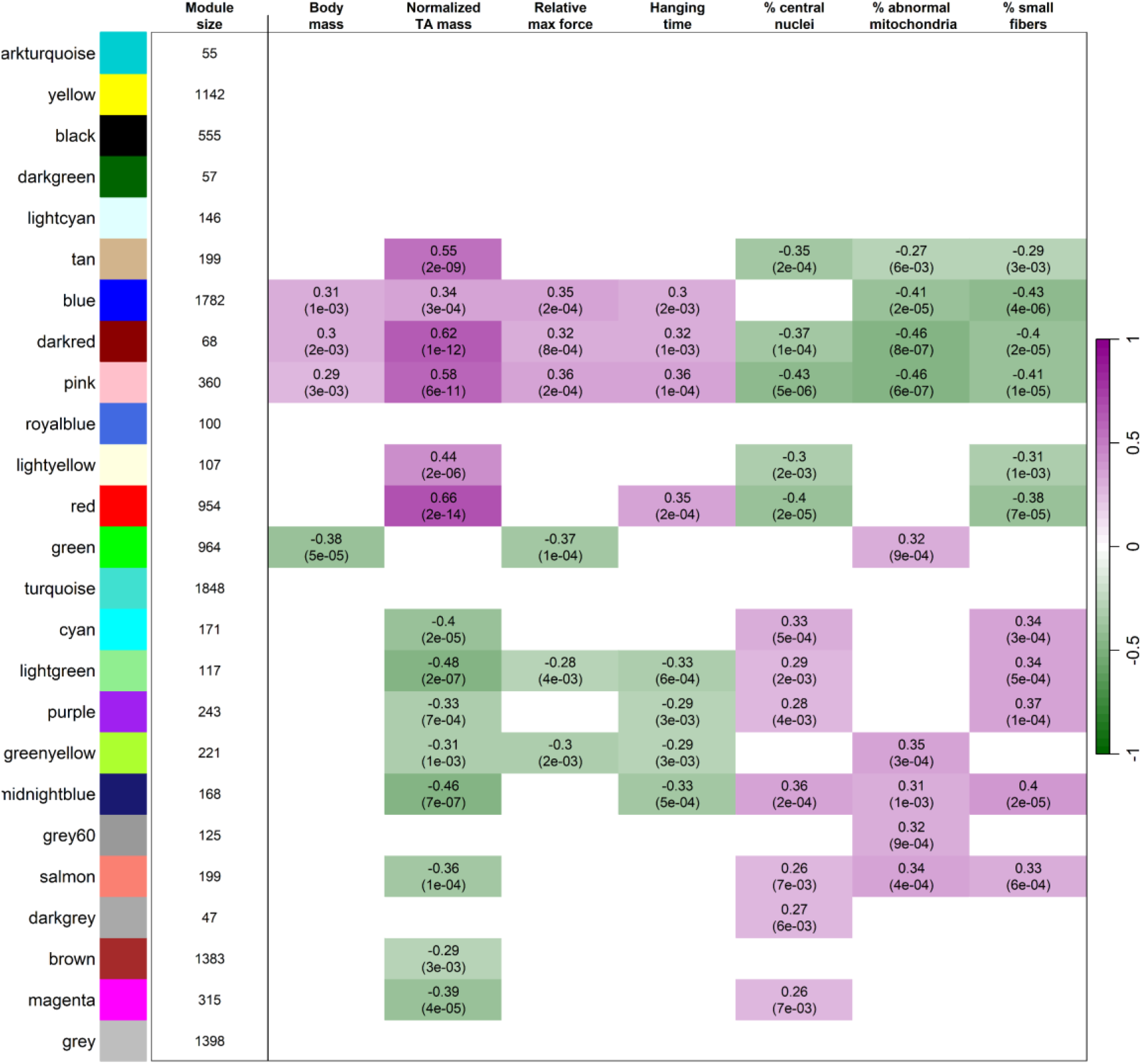
WGCNA module-trait correlation heatmap. Rows correspond to modules (designated by color names) and columns to phenotypic traits. The “grey” module contains genes unassigned to any module. Purple cells indicate a positive correlation between gene expression within a module and quantitative phenotypic traits, while green cells represent a negative correlation. Only statistically significant correlations (Bonferroni-adjusted *P* value < 0.05) are shown. For each significant module-trait pair, the Pearson correlation coefficient and Bonferroni-adjusted *P-*value (in parenthesis) are indicated.

#### Enrichment analysis of modules significantly correlated to phenotypic traits

To identify the biological functions associated with each module, Gene Ontology (GO) terms over-representation analysis was conducted for the biological process (BP), cellular component (CC), and molecular function (MF) ontologies (Fig. 3, Supplementary Information S2). Among the beneficial modules, the blue module was enriched in genes associated with apoptosis regulation, calcium ion transport and muscle development and contraction (Fig. 3A). The pink module was enriched in genes associated with RNA processing, transcription, and translation, and the dark red module was associated with oxidative phosphorylation (Fig. 3B). Among pathogenic modules, the brown module was associated with immune system processes. The midnight blue, salmon, light green and grey60 modules were associated with vascularization and/or innervation, with enriched pathways related to angiogenesis and with the development, function and maintenance of the neuromuscular junction (axonogenesis, synaptic transmission, cell-cell junction, semaphoring-plexin signaling). The green yellow module was associated with beta oxidation of fatty acids (Fig. 3C), and the purple module was associated with histone deacetylation and autophagy (Fig. 3D). Overall, these findings underscore multiple pathways linked to muscle structure and function across three murine models of CNM, as well as their modulation through therapeutic intervention.

**Fig. 3.**
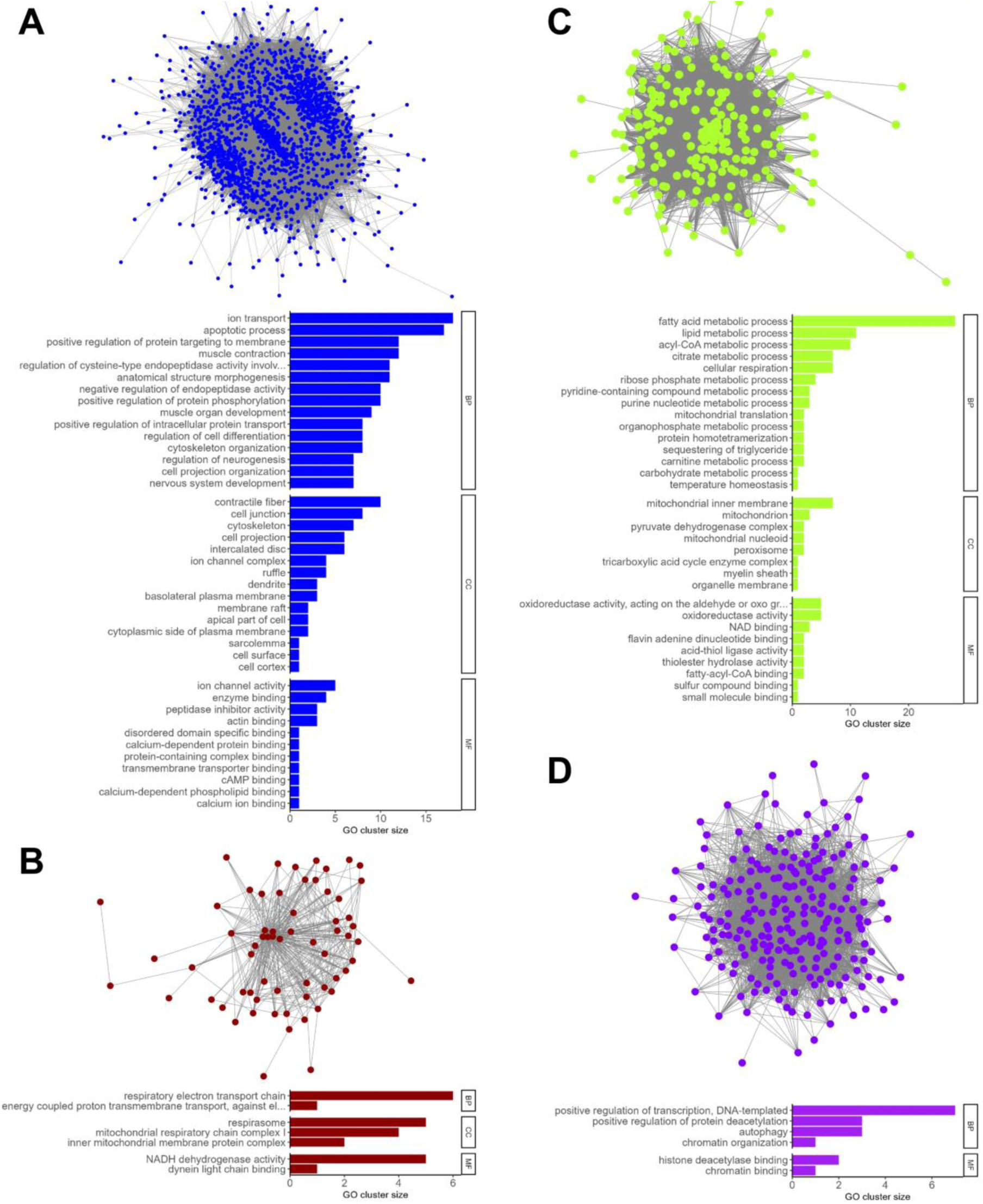
Visualization and GO term over-representation analysis of selected WGCNA modules. **(A)** Blue module **(B)** Dark red module **(C)** Green yellow module **(D)** Purple module

### Multi-omics integration in X-linked CNM

To investigate the mechanisms implicated in the pathology of XLCNM, we integrated experimental omics data (transcriptomics, proteomics, metabolomics) obtained from *Mtm1*^-/y^ mice and their WT littermates with public databases. To integrate these diverse and multiscale data sources, we created a multilayer network composed of a gene multiplex network, a metabolite multiplex network, and three monoplex networks containing phenotypes, metabolic reactions and tissues (Fig. 4). These networks are connected to each other by bipartite networks representing several types of relationships, such as localization, expression, pathway co-membership, or participation in a reaction described in the murine genome-wide metabolic model (Table 2, Table 3).

**Fig. 4.**
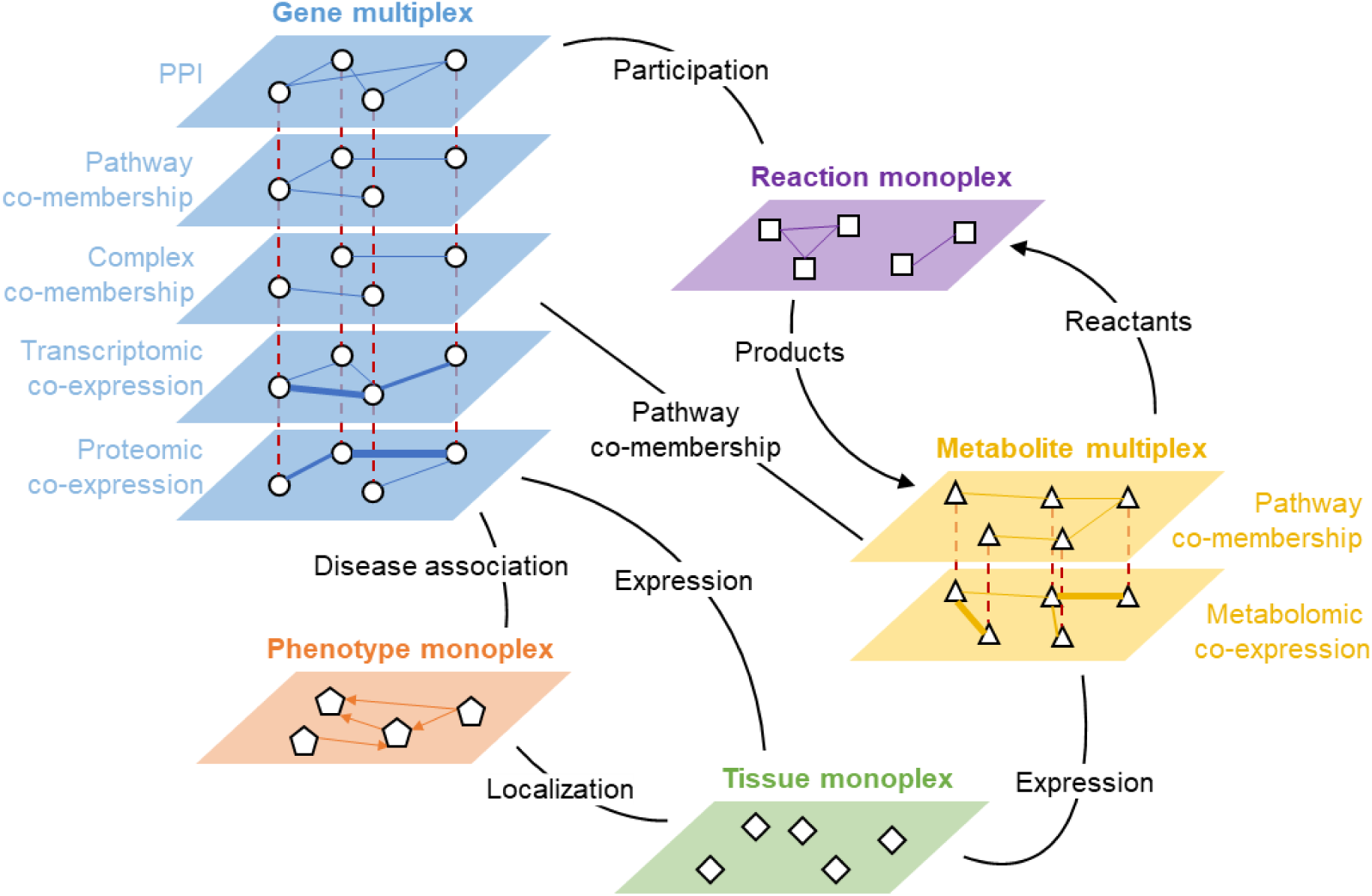
Schematic representation of the multilayer network for X-linked CNM. Each multiplex network contains a single type of nodes that are connected to each other within each layer (continuous lines) and to themselves across layers (red dashed lines). The bipartite networks linking the multiplex networks to each other are schematized as black lines. Arrows indicate edge directionality and line width indicates edge weight. PPI: protein-protein interaction.

**Table 2.**
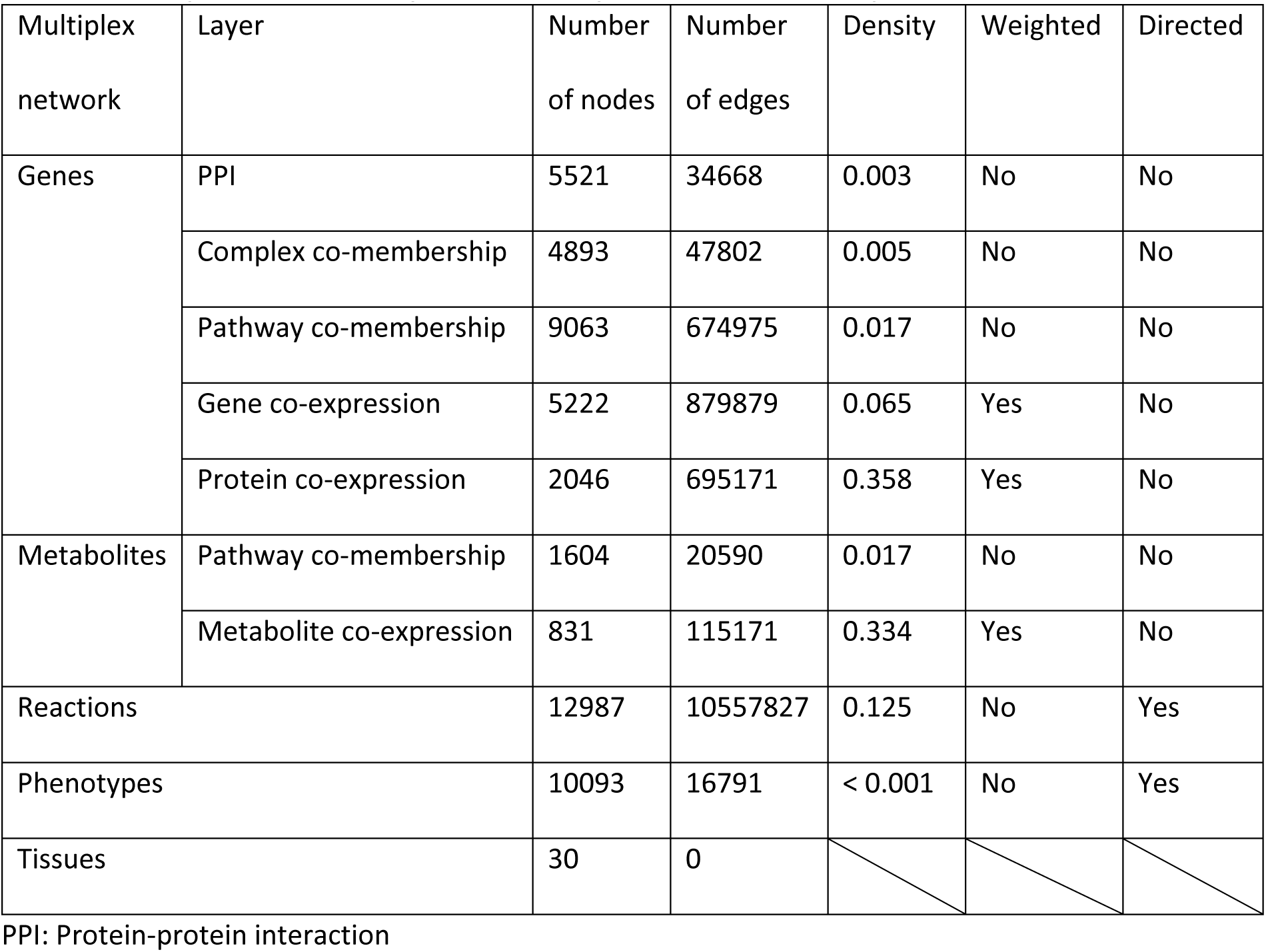
Description of the multiplex networks present in the multilayer network.

| Multiplex network | Layer | Number of nodes | Number of edges | Density | Weighted | Directed |
| --- | --- | --- | --- | --- | --- | --- |
| Genes | PPI | 5521 | 34668 | 0.003 | No | No |
|  | Complex co-membership | 4893 | 47802 | 0.005 | No | No |
|  | Pathway co-membership | 9063 | 674975 | 0.017 | No | No |
|  | Gene co-expression | 5222 | 879879 | 0.065 | Yes | No |
|  | Protein co-expression | 2046 | 695171 | 0.358 | Yes | No |
| Metabolites | Pathway co-membership | 1604 | 20590 | 0.017 | No | No |
|  | Metabolite co-expression | 831 | 115171 | 0.334 | Yes | No |
| Reactions |  | 12987 | 10557827 | 0.125 | No | Yes |
| Phenotypes |  | 10093 | 16791 | < 0.001 | No | Yes |
| Tissues |  | 30 | 0 |  |  |  |

PPI: Protein-protein interaction

**Table 3.** Description of the bipartite networks present in the multilayer network.

| Bipartite network | Number of nodes | Number of edges | Density | Weighted | Directed |
| --- | --- | --- | --- | --- | --- |
| Genes - metabolites | 6062 | 69470 | 0.004 | No | No |
| Genes - reactions | 10115 | 22788 | 0.004 | No | No |
| Metabolites - reactions | 15047 | 18867 | < 0.001 | No | Yes |
| Reactions - metabolites | 13460 | 18064 | < 0.001 | No | Yes |
| Genes - tissues | 16683 | 533921 | 0.002 | Yes | No |
| Genes - phenotypes | 15134 | 262415 | 0.002 | No | No |
| Metabolites - tissues | 3384 | 6425 | 0.001 | No | No |
| Phenotypes - tissues | 3926 | 4569 | 0.001 | No | No |

To explore the network, random walk with restart (RWR) was performed using the Python package MultiXrank^13^. This approach simulates a random particle navigating the network. At each step, the particle can either move to a neighboring node or restart its walk from a node that is randomly selected from a user-provided set of seed node(s). RWR output scores were used to rank and identify top-scoring nodes that are closely connected to the seed(s).

#### Identification and removal of nonspecific hub nodes with random seeding

After performing RWR on the full multilayer network with several sets of seeds, we observed that generic chemicals such as water, H^+^, or ATP were often present in the list of top-scoring metabolites. We hypothesized that these highly connected nodes, or "hubs," could create artificially shortened paths and be inaccurately ranked among the top-scoring nodes by RWR. To assess the presence of hubs in the multilayer network, we conducted RWR using 1000 random seeds, each representing a randomly selected protein-coding gene. To assess the impact of including experimental data on hubs, RWR was performed on both the full multilayer network and a subset excluding the experimental layers based on transcriptomic, proteomic, and metabolomic data (called “subset” hereafter). For each RWR iteration, the top-scoring genes and metabolites were identified, and the number of occurrences of each node among the top 10, 50 or 100 ranked nodes was calculated (**Fig. 5**).

**Fig. 5.**
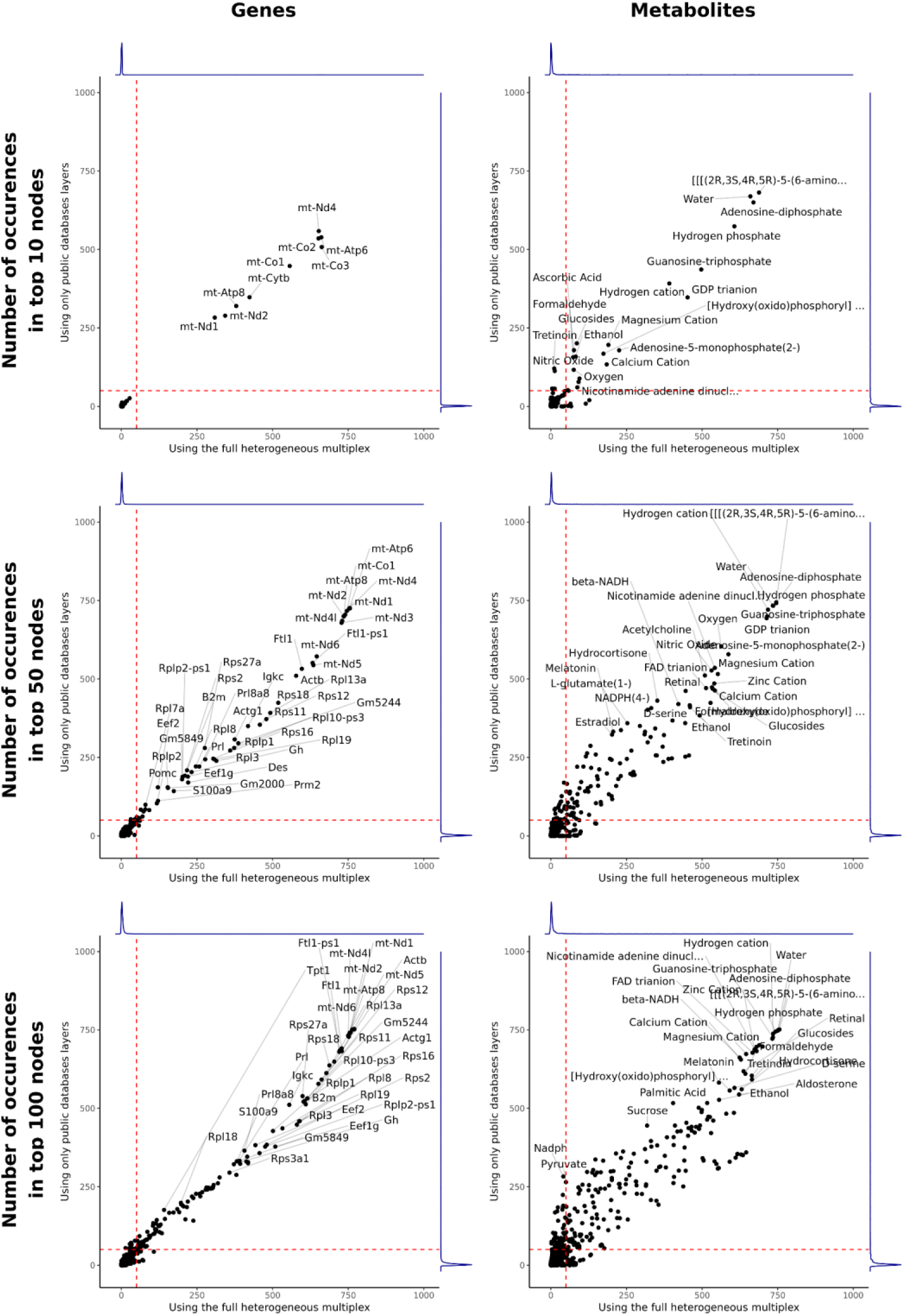
Number of occurrences in the top scoring nodes for 1000 RWR with random seeds. 1000 seeds were created by sampling a random protein-coding gene of the mouse genome. For each random seed, RWR was carried out on the full multilayer network and on a subset of the network without experimental layers. The lists of the top 10, 50, and 100 scoring genes, metabolites and phenotypes were determined for each run. Nodes are plotted according to the number of times they occur in the top-scoring lists, with the full multilayer network runs on the x-axis and the multilayer subset runs on the y-axis. Red dashed lines indicate 5% of the total number of runs. Blue lines show the densities of points along each axis.

When considering the top 10 genes, we observed an over-representation of mitochondrial genes using both the full multilayer network and its subset, with 250 to 750 occurrences in the top nodes out of 1000 iterations. Additionally, we observed an over-representation of water, energy-related molecules (ATP, ADP, GTP, and GDP), and several cations (H^+^, Ca^2+^, Mg^2+^) in the top 10 scoring metabolites. When considering the lists of top 50 or top 100 scoring nodes, we observed an increased number of hubs, including genes encoding ribosomal protein large and small subunits, and additional energy-related metabolites such as NADH and FAD. A higher correlation between full multilayer and subset runs can be observed for the genes compared to the metabolites. This indicates that the metabolomic co-expression layer has a higher impact on the network than the transcriptomic and proteomic layers.

This may be due to the difference in composition between the gene multiplex network (three layers extracted from public databases, two experimental layers) and the metabolite multiplex network (one layer extracted from a public database, one experimental layer).

Nonetheless, when considering the top 10, 50, or 100 nodes, density plots showed that the vast majority of top-scoring nodes occurred in less than 5% of iterations. Therefore, we considered that genes and metabolites ranked in the top 50 scoring nodes in more than 5% of iterations were nonspecific hub nodes and represented false-positive hits. For the following analyses, nonspecific hub nodes were removed from the multilayer network.

#### Investigation of the pathomechanisms of XLCNM using a multilayer heterogeneous network

After removing nonspecific hub nodes from the multilayer network, we explored it using RWR in order to investigate the pathomechanisms implicated in XLCNM. To do this, we performed RWR with several sets of seeds and selected the top 20 highest-scoring gene and metabolite nodes.

We sought to identify nodes that may be implicated in several phenotypes observed in XLCNM, such as skeletal muscle atrophy (HP:0003202), muscle weakness (HP:0001324), and centrally nucleated skeletal muscle fibers (HP:0003687). Therefore, we conducted RWR on the multilayer network, using each of these phenotypes and *Mtm1* as joint seeds. We observed a strong overlap among the 20 top-scoring metabolites for these three sets of seeds. Shared hits included inositol (Ins), phosphatidylinositol (PtdIns) and phosphoinostide (PtdIns*P*) related metabolites: Ins, PtdIns pool, PtdIns(4)*P*, PtdIns(3,4)*P*_2_, Ins(1,3,4)*P*_3_, Ins(1,3,4,6)*P*_4,_ Ins(1,3,4,5)*P*_4_ and scyllo-Ins(4)*P*. These metabolites are directly linked to the function of *Mtm1*, which encodes myotubularin, a phosphoinositides phosphatase. Other shared top metabolites were ADP, lutein, geranylgeranyl diphosphate (GGPP), arachidonate, and S-adenosylmethionine. By comparison, the overlap was smaller in the 20 top-scoring genes, with only *Mb* and *Myh7* (which are both highly expressed in slow-twitch oxidative fibers) as common hits for the three seeds. Additionally, *Mtmr12* (encoding a myotubularin-related protein that binds MTM1)*, Ckm* (encoding a cytoplasmic enzyme involved in energy homeostasis)*, Myl2* (encoding a sarcomeric protein expressed in cardiac muscle and slow-twitch skeletal muscle fibers)*, Pten* (encoding a PtdIns(3,4,5)*P*_3_ phosphatase) and *mt-Te* (a mitochondrially encoded tRNA) were common hits for the muscle weakness and skeletal muscle atrophy seeds. Seed-specific top-scoring nodes are detailed in Table 4. Specific top scoring genes for the three sets of seeds are all implicated in neuromuscular disorders described in the Muscle Gene Table^14^ except *Pik3r5,* a regulatory subunit of phosphatidylinositol 3-kinases that phosphorylate PtdIns to PtdIns3*P*, *Mtmr14* that encodes a myotubularin-related protein, and *Myf6*, a myogenic factor. Overall, using a multilayer heterogeneous network and RWR analysis, this study identified key metabolites and genes linked to *Mtm1* function and XLCNM phenotypes, shedding light on shared and phenotype-specific molecular mechanisms.

**Table 4.** Specific top scoring genes and metabolites using *Mtm1* and several XLCNM phenotypes as seeds.

| Seeds | Genes | Metabolites |
| --- | --- | --- |
| *Mtm1*<br>HP:0003202<br>(skeletal muscle atrophy) | *Trpv4*<br>*Capn3*<br>*Lmna*<br>*Neb*<br>*Ryr1*<br>*Pomt2*<br>*Fkrp*<br>*Crppa*<br>*Tnnt1*<br>*Fktn*<br>*Inpp5k*<br>*Pomt1* | Dexamethasone<br>Diacetyl<br>Epitestosterone<br>Oxytocin |
| *Mtm1*<br>HP:0001324 (muscle weakness) | *Fhl1*<br>*Tnnc2*<br>*Klhl41*<br>*Chrna1*<br>*Itga7*<br>*Col6a1*<br>*Col6a2*<br>*Scn4a*<br>*Col12a1*<br>*Hnrnpa2b1*<br>*Col13a1*<br>*Pik3r5* | 9-O-acetylneuraminic acid<br>Einecs 307-780-4<br>(2s)-1-(Alpha-D-Glucopyranosyloxy)-3-(Hexadecanoyloxy)propan-2-yl (11z)-Octadec-11-Enoate<br>1-O-(alpha-D-galactopyranuronosyl)-N-tetradecanoyldihydrosphingosine |
| *Mtm1*<br>HP:0003687 (centrally nucleated skeletal muscle fibers) | *Mtmr14*<br>*Myot*<br>*Hnrnpa1l2-ps2*<br>*Hnrnpa1*<br>*Selenon*<br>*Pyroxd1*<br>*Adgrg6*<br>*Bves*<br>*Ccdc78*<br>*Syne1*<br>*Sil1*<br>*Matr3*<br>*Unc45b*<br>*Fxr1*<br>*Orai1*<br>*Tpm3*<br>*Myf6* | Thromboxane A2<br>Tetrahydrobiopterin<br>Uric Acid<br>9-Anthroic acid<br>Naadp<br>Niflumic acid<br>Aspirin<br>Diacetyl |

## Discussion

This study provides a comprehensive molecular perspective common to several forms of centronuclear myopathies by integrating transcriptomic data with phenotypic profiles obtained from several murine models of CNM treated with various therapeutic approaches. XLCNM was further investigated by integrating multi-omics data with publicly available biological knowledge. These findings contribute to a more detailed understanding of the molecular landscape of CNM and suggest new directions for targeted therapeutic interventions. Limitations of our study include the use of average group values for phenotypic traits instead of individual data points, which may mask subtle within-group variations. Additionally, while murine models provide valuable insights, they may not fully recapitulate human CNM pathology, especially concerning the distribution of muscle fiber types and genetic background diversity. Future studies should seek to verify these findings in human tissues or *in vitro* models to confirm the translatability of our results.

### Gene co-expression modules and phenotype correlation

Employing WGCNA facilitated the identification of gene modules linked to CNM phenotypes, highlighting both pathogenic and beneficial molecular processes relevant to muscle function and structure. Notably, beneficial modules were enriched for genes involved in apoptosis regulation, calcium ion transport and muscle contraction, oxidative phosphorylation, and RNA processing, which are essential for maintaining muscle cell integrity and function. The enrichment of genes in these modules may underpin molecular mechanisms promoting muscle resilience in response to CNM-related stressors. Conversely, pathogenic modules were linked to immune system processes and structural disruptions in the muscle tissue, such as defects in vascularization and innervation. These insights align with recent findings of immune cells infiltration in CNM-affected murine muscle^6^ and reports of neuromuscular junction defects in CNM patients and animal models^15–17^. However, vascularization defects have yet to be reported in CNM and may represent an intriguing area for future research to uncover their potential role in disease pathology. In terms of energy metabolism, one pathogenic module was enriched in genes linked to fatty acid oxidation. Excessive fatty acid oxidation has been linked to skeletal muscle atrophy in cancer cachexia, and dietary supplementation with nicotinamide or lipids has been shown to improve muscle health in this context^18,19^. This finding suggests the possibility of using dietary interventions, such as high-fat diets, as adjunctive therapies to augment muscular resilience in CNM.

### Implications of multi-omics integration and network exploration

Our integration of multi-omic data with public databases through a multilayer network enabled a high-resolution view of XLCNM pathomechanisms. The multilayer network, comprising gene, metabolite, reaction, phenotype and tissue multiplex networks, facilitated the ranking of key metabolites and genes connected to XLCNM phenotypes, including muscle atrophy, muscle weakness, and abnormal nuclear centralization in muscle fibers. The presence of *Myh7* and *Mb* (which are highly expressed in oxidative fibers) in the top-ranking genes suggests a heightened role of oxidative metabolism in XLCNM pathology, aligning with the WGCNA results on several CNM models discussed above. The emergence of inositol and phosphoinositide metabolites as top-ranked hits across multiple XLCNM phenotypes is consistent with the function of myotubularin, the protein encoded by *Mtm1*, but provides limited new insights into the disease. On the other hand, other top-scoring metabolites may provide novel therapeutic opportunities to improve muscle phenotype. Indeed, dietary lutein has been associated to a reduction in oxidative stress and inflammation in rat, increased muscle fiber diameter in chicken, and may act through regulation of *MSTN*, a gene that is downregulated in several CNM models^6,20,21^. GGPP depletion has also been shown to regulate *MSTN* and cause skeletal muscle atrophy in mice, and mutations in *GGPS1*, encoding a GGPP synthase, have been linked to congenital myopathies^22,23^. Additionally, GGPP depletion has been linked to apoptosis and the reduction of the expression, localization, and GTP-binding ability of the Rho-GTPase RAC1 in cell studies^24^. RAC1 plays a role in recruiting MTM1 to plasma membrane ruffles, membrane protrusions implicated in actin cytoskeleton remodeling ^25^. Finally, S-adenosylmethionine increase through betain-supplemented diet has been shown to delay age-related muscle loss by promoting protein synthesis^26^.

## Conclusion

In summary, this study leveraged multi-omic integration and network analysis to unravel key molecular drivers of CNM pathology. The findings enhance our understanding of CNM pathomechanisms, especially concerning structural disruptions and metabolic regulation. The insights gained here could inform the development of novel therapeutic strategies, including dietary interventions, to mitigate the clinical signs of CNM. Future research should aim to refine these molecular targets and evaluate potential treatments in translational models to advance therapeutic options for CNM patients.

## Materials and methods

### Transcriptomic data

Bulk RNA-sequencing datasets from various mouse models were retrieved from previously published studies and are available online at NCBI’s GEO through the GEO Series accession number GSE160084^6^. Additionally, the transcriptomic characterization of tamoxifen-treated *Dnm2*^S619L/+^ and *Bin1*^mck-/-^ mice described in ^7^ have been deposited on NCBI’s GEO and are accessible through GEO Series accession number GSE282489. All studies were performed in tibialis anterior (TA) muscle. Detailed information about each cohort is available in Table 1.

### Proteomic data

Proteomic profiling of cohort A was retrieved from a previous study and the data are available in the PRIDE repository with the identifier PXD021765 ^6^.

### Metabolomic data

Metabolomic profiling of quadriceps muscles from 5 week-old *Mtm1*^-/y^ and WT mice was carried out by Metabolon (NC, USA). Briefly, 50-130mg of muscle and 150-300μL of serum were flash-frozen and sent to Metabolon. Samples were prepared using the automated MicroLab STAR® system from Hamilton Company. Ultrahigh-performance liquid chromatography–tandem mass spectroscopy (UPLC–MS/MS) was carried out with a Waters ACQUITY ultra-performance liquid chromatography (UPLC) and a Thermo Scientific Q-Exactive high resolution/accurate mass spectrometer interfaced with a heated electrospray ionization (HESI-II) source and Orbitrap mass analyzer operated at 35,000 mass resolution. Raw data were extracted, peak-identified and QC processed using Metabolon’s hardware and software. Peaks were quantified using area-under-the-curve.

### Weighed gene correlation network analysis

WGCNA was carried out using the bulk RNA-seq data from the 7 cohorts (Table 1). To construct the weighed gene correlation networks, genes with more than 10 reads in at least 90% of all samples were considered. Gene expression normalization was performed with DESeq2^27^. The gene correlation network was built with the WGCNA R package ^4^. Briefly, the adjacency matrices were computed with a soft power value of 12. Topological overlap dissimilarity matrices were computed and used to perform hierarchical clustering of the genes. Gene modules were determined with dynamic tree cut and close modules (dissimilarity < 0.25) were merged. Module-trait Pearson correlations were computed, the associated *P*-values were determined with Student’s t-test and adjusted with Bonferroni correction based on the number of traits.

### Multi-omics integration with knowledge from public databases

#### Biological network construction

The composition of the multiplex-heterogeneous biological network used to integrate transcriptomic, proteomic and metabolomic data with publicly available knowledge is detailed below. The full network contains five multiplex networks (one or more layers connecting one type of node) and bipartite networks encoding links between the different multiplex networks. Unless specified otherwise, the layers are unweighted and undirected.

Description of the multiplex networks (Table 2):

- A gene multiplex network composed of five layers encoding different gene-gene interactions. Three layers were constructed based on publicly available knowledge: protein-protein interactions (PPI) extracted from the STRING database (experimental confidence score > 700), molecular complex co-membership extracted from the CORUM database, and pathway co-membership extracted from the Reactome database^28–30^. The remaining two weighted layers were constructed with WGCNA of bulk transcriptomic and proteomic datasets from WT and *Mtm1*^-/y^ mice. For the transcriptomic layer, samples of cohorts (A-C) were used. After gene expression normalization and batch effect correction with DESeq2 and limma^31^, the topological overlap matrix was computed using a soft power of 10, and edges with an adjacency > 0.15 were exported. The proteomic layer was built similarly. The wrMisc and wrProteo R packages were used to normalize and to impute missing data with default parameters. Then, the topological overlap matrix was computed using a soft power of 12, and edges with an adjacency > 0.25 were exported.
- A metabolite multiplex network composed of two layers: one layer encoding pathway co-membership extracted from the Reactome database and one weighted layer built with WGCNA from metabolomics data (WT and *Mtm1*^-/y^ samples, soft power = 5, adjacency > 0.05)
- A directed monoplex network containing phenotypes described in the Human Phenotype Ontology (HPO)^32^. Directed edges linking HPO terms to their parents were retrieved using custom SPARQL queries on the OWL file provided by the HPO and loaded in Apache Foundation’s JENA suite (v4.7.0) (Supplementary Information S1).
- A monoplex network containing the reactions described in the mouse genome-scale metabolic model (Mouse-GEM)^33^. Reactions belonging to the same subsystem (sharing a similar metabolic function) are linked together.
- A monoplex network containing tissues found in the Genotype-Tissue Expression project (GTEx)^34^ and linked to other multiplexes through bipartite networks.

Description of the bipartite networks (Table 3):

- A weighted gene-tissue bipartite network extracted from the GTEx project. This bipartite network links genes to the tissues in which they are expressed. Median transcript per million (TPM) values were used as edge weights, only keeping edges with TPM>5.
- A gene-metabolite bipartite network extracted from Reactome based on pathway co-membership^30^.
- A metabolite-tissue bipartite network extracted from the Human Metabolome Database (HMDB) linking metabolites to the tissues in which they have been detected^35^.
- Three bipartite networks extracted from the Mouse-GEM: two directed networks linking metabolites to reactions (reactants to reaction and reaction to products), and one network linking genes to the reactions in which the proteins they encode are implicated^33^.
- Two bipartite networks extracted from the HPO: one network linking phenotypes to the genes associated to diseases in which the phenotype can be observed, and one network linking phenotypes to the tissues they affect. The second network was obtained using custom SPARQL queries on the OWL file provided by the HPO and loaded in Apache Foundation’s JENA suite (v4.7.0) (Supplementary Information S1).

#### Heterogeneous multiplex exploration

The heterogeneous multiplex was explored with random walk with restart (RWR) using the Python package MultiXrank^13^ with global restart probability r = 0.7 and uniform inter-layers jump, layer restart, inter-multiplex networks jump, and multiplex network restart probabilities. In practice, this means that all layers had the same weight within each multiplex network, and that all multiplex networks had the same weight within the multilayer network.

### Metabolite ID mapping

To combine experimental metabolomic data, metabolite information extracted from the Mouse-GEM, Reactome, and HMDB, metabolites were mapped to their PubChem ID if possible. Metabolite ID mapping was performed programmatically by sending custom REST queries to the PubChem Identifier Exchange Service using cross reference IDs as input. For the metabolomics data, mapping was performed based on cross-references (PubChem, HMDB, KEGG) and INCHI keys provided by Metabolon. For the Mouse-GEM data, cross-references provided with the GEM were combined with RECON3D^36^ and MetaNetX^37^ information to obtain ChEMBL, BioCyc, ChemSpider, KEGG, HMDB, LipidMaps, ChEBI cross-reference IDs and map them to PubChem IDs. If ID mapping was unsuccessful, original IDs were kept. For the Reactome database, mapping was performed on ChEBI IDs, and unmapped metabolites were discarded.

### Network visualization

Cytoscape was used to visualize networks and sub-networks^38^.

### Enrichment analyses

GO term enrichment analysis was performed with the R package ClusterProfiler, and enriched GO terms were clustered based on their semantic similarity^39,40^.

## Acknowledgements

The work of the Interdisciplinary Thematic Institute IMCBio, as part of the ITI 2021-2028 program of the University of Strasbourg, CNRS and Inserm, was supported by IdEx Unistra (ANR-10-IDEX-0002), and by SFRI-STRAT’US project (ANR-20-SFRI-0012) and EUR IMCBio (ANR-17-EURE-0023) under the framework of the France 2030 Program. This work was also funded by Association Française contre les Myopathies-Téléthon (23933), the Muscular Dystrophy Association (576154), and the Fondation pour la Recherche Médicale (201903007992). AS is a fellow of Ecole Inserm Pfizer Innovation and recipient of the Société Française de Myologie Master research fund.

## Authors contributions

AS: Conceptualization, Data curation, Formal Analysis, Funding acquisition, Investigation, Methodology, Software, Visualization, Writing – original draft; CG: Investigation, Resources; DR: Investigation, Resources; JT: Supervision, Writing – original draft; JL: Conceptualization, Funding acquisition, Supervision, Writing – original draft

## Supplementary information captions

Supplementary information S1. Custom SPARQL queries used to process the HPO database.

Supplementary Information S2. Phenotypic data used for WGCNA module-trait correlation.

Supplementary Information S3. Full enrichment analysis results of WGCNA modules.

## Data availability statement

All relevant data are within the paper and its Supplementary Information files. The Supplementary Information Files and the code used to produce the results were deposited on Zenodo at https://doi.org/10.5281/zenodo.14225124.

## Notes

https://zenodo.org/records/14225124

### Competing Interest Statement

The authors have declared no competing interest.

